# Selective activation of estrogen receptors modulates electrical activity and insulin secretion in human EndoC-βH1 cells

**DOI:** 10.64898/2026.11.10.763916

**Authors:** Roberto Sempere-Navarro, Tania Exposito, Cristina Quereda, Marina Serrano-Macia, Ángel Nadal, Juan Martinez-Pinna

## Abstract

Estrogen receptors (ERs), ERα and ERβ have important roles in the physiology of the endocrine pancreas in animal models but the effects of ERs activation on insulin-secreting human cells are still poorly understood. Here, we investigated the role of ERα and ERβ as modulators of electrical activity and insulin secretion in the insulin-secreting human cell line EndoC-βH1. Immunocytochemical analysis revealed a widespread cellular distribution of ERα, while ERβ was predominantly localized outside the nucleus. We observed that both, estradiol (E2) and the ERβ agonist diarylpropionitrile (DPN), led to a decrease in insulin secretion in the absence of stimulatory glucose concentrations, without affecting insulin content. Patch clamp experiments revealed that a 48 hours treatment with either the ERα agonist PPT or DPN resulted in depolarization of the resting membrane potential, increased action potential firing frequency, and altered the action potential waveform in EndoC-βH1 cells. PPT increased voltage-dependent Ca²⁺ channel currents, while DPN reduced K⁺ and Na⁺ currents. These results suggest that activation of ERα and ERβ modulate β-cell electrical activity through distinct pathways on ion channels. The actions of E2, PPT and DPN were not caused by altered expression levels of ion channel genes after 48 hours of treatment. This suggests that these receptors act extra-nuclearly without affecting gene expression. Together, our findings demonstrate that selective activation of ERα and ERβ modulate β-cell excitability through distinct electrophysiological mechanisms and that E2 and DPN reduced basal insulin secretion in human EndoC-βH1 cells. These results support ERα and ERβ as potential targets for modulating human pancreatic β-cell function.

## Introduction

Pancreatic β-cells synthesize, store and release insulin, which is essential for maintaining blood glucose levels within physiological limits. Decreased insulin release in response to glucose results in chronic hyperglycemia, a hallmark of type 2 diabetes (T2D). This metabolic disease affects over 589 million people around the world, a figure that is projected to rise to 853 million by 2050 (1). Therefore, it is important to understand the physiological mechanisms that may improve pancreatic β-cell function as well as β-cell survival.

Estrogens have been demonstrated to regulate β-cell function (2–4) and to protect β-cells from death-induced insults in rodent models (5, 6). During pregnancy, estrogens may help the pancreas to adapt to increased metabolic demands and peripheral insulin resistance. Failure to do so may result in gestational diabetes (7, 8). Estrogens may also play a role in sex differences. For example, they may explain, at least in part, why women during fertile age have lower rates of diabetes than men, though this trend reverses after menopause (9, 10).

Estrogens exert their biological actions through estrogen receptors, ERα and ERβ (11, 12) activating nuclear-initiated and extranuclear-initiated pathways (13). Extranuclear-initiated pathways also involve the G-protein–coupled estrogen receptor (GPER) (14, 15).

Both, ERα and ERβ regulate stimulus-secretion coupling in mouse pancreatic β-cells. ERα regulates insulin gene expression and content via an extranuclear-initiated pathway involving the activation of ERK1/2 and the transcription factor NeuroD1 (3, 16). ERα also protects β-cells from apoptosis-inducing agents (17, 18). ERβ decreases K_ATP_ channel activity within minutes and rapidly increases glucose-stimulated insulin secretion (GSIS) through an extranuclear-initiated pathway involving cGMP and membrane guanylate cyclase (19). ERβ agonists act as enhancers of GSIS and improve glucose homeostasis in rodent models of diabetes (4) and regulate ion channel expression and electrical activity (20). The role of GPER is less studied but is involved in rapid regulation of GSIS by estradiol (21, 22) and in the protection of β-cell death (23).

In the present study, we examined the effects of ERα and ERβ agonists on the stimulus secretion coupling in the insulin-secreting human cell line EndoC-βH1. The limited availability of human islets together with donor variability, has made studying human β-cells difficult. The EndoC-βH1 human β-cell line is increasingly being used to study various aspects of human β-cell biology (24–26). Importantly, stimulus-secretion coupling, including the set of ion channels responsible for the electrical activity, is similar to that of human β-cells (24). Here, we demonstrate that agonists of ERα and ERβ decrease basal insulin secretion, alter ion channel currents and modified action potential firing in EndoC-βH1 cells after 48 hours of exposure. These effects were not accompanied by changes in the transcription of the main ion channel genes.

Our results demonstrate that estrogen receptor agonists have modulatory actions in insulin secretion on human EndoC-βH1 cells, suggesting their potential use in diabetes treatment.

## Material and methods

### Chemicals

The estrogen receptor α-selective agonist propylpyrazoletriol (PPT; Cat. No. H6036), the estrogen receptor β-selective agonist diarylpropionitrile (DPN; Cat. No. H5915), and 17β-estradiol (E2; Cat. No. E8875) were purchased from Sigma-Aldrich (St. Louis, MO, USA). All chemicals were dissolved in dimethyl sulfoxide (DMSO) to prepare the stock solution. The amount of DMSO was maintained at less than 0.03%.

### Cell Culture

EndoC-βH1 cells (Univercell Biosolutions, Paris, France) were cultured and passaged as previously described [79]. Briefly, cells were cultured in DMEM containing 5.6 mM glucose, 2% BSA fraction V fatty acid free (Sigma-Aldrich, Saint Louis, MO, USA), 50 μM 2-β-mercaptoethanol (Gibco, Paisley, UK), 10 mM nicotinamide, 5.5 μg/mL human transferrin, 6.7 ng/mL sodium selenite (Sigma-Aldrich, Saint Louis, MO, USA), 100 units/mL penicillin and 100 μg/mL streptomycin (Thermo Fischer Scientific, Waltham, MA, USA). Cells were seeded at a density of 70–80 × 103 cells/cm^2^ on ECM (1%) and fibronectin (2 μg/mL) (Sigma-Aldrich) -coated plates, and cultured at 37 °C in 5% CO2. For treatment with estrogenic agonists, DMEM was replaced with DMEM without phenol red (Sigma-Aldrich) and BSA was replaced by 2% charcoal dextran-treated FBS (HyClone, GE Healthcare Life Sciences, Logan, UT, USA). EndoC-βH1 cells were incubated at 37 °C with 5% CO^2^. Cells were discarded after passage 72.

### Immunofluorescence

A total of 200 × 10^3^ EndoC-βH1 cells per well were seeded onto coated 24-well plates and cultured at least 24 h at 37 °C with 5% CO_2_. Subsequently, EndoC-βH1 cells were rinsed very carefully so as not to detach with PBS, fixed with 4% paraformaldehyde dissolved in PBS for 15 min, washed three times with PBS, permeabilized with 0.2% Triton X-100 diluted in PBS for 15 min and washed twice with PBS. To block non-specific antibody binding, samples were incubated with 1% BSA in PBS for 1 h. in a humid chamber. Primary antibodies were purchased from Invitrogen, Thermo Fischer Scientific, Waltham, MA, USA, against ERα (Cat. MA5-14501, 1:50), ERβ (Cat. MA5-24807, 1:200), KDEL (Cat. PA1-013, 1:100), GM130 (Cat. PA5-85643, 1:100), and cytochrome c (Cat. PA5-87380, 1:100) were diluted in blocking solution and applied overnight at 4 °C in a humidified chamber. After washing three times, the samples were incubated for 1 h in a humidified, light-protected chamber, in the presence of a 1:500 dilution of goat secondary antibodies against rabbit (Cat. A-21428; Cat. A-32731) or mouse (Cat. A-32727) IgG, conjugated to either Alexa Fluor 488 or 555. Then the samples were incubated with Hoechst 33342 (Invitrogen, H3570) diluted 1:1000 in PBS for 10 min to visualize the nuclei. Negative-control samples were subjected to the same immunocytochemical procedure as the experimental samples, except that the primary antibody was omitted. Fluorescence was detected using a Zeiss® LSM 900 confocal laser-scanning microscope with Airyscan 2 (Carl Zeiss Microscopy GmbH, Jena, Germany).

The JACoP plugin, which was developed for the NIH ImageJ v1.52p software, was used to assess the colocalization. The degree of colocalization was evaluated using the Manders colocalization coefficient (MCC), with values near 1 indicating complete colocalization and those near 0 indicating absence of colocalization (27, 28). In some studies, the colocalization ranges are defined as follows: very high, 0.8 ≤ MCC ≤ 1.0; high, 0.6 ≤ MCC < 0.8; moderate, 0.4 ≤ MCC < 0.6; low, 0.2 ≤ MCC < 0.4; and not detectable, 0 ≤ MCC < 0.2 (29). MCC values were expressed as the mean ± SD from 6 cells analyzed from micrographs obtained in triplicate experiments, in which the cells of interest were cropped using the NIH ImageJ software before analysis.

### Insulin Secretion and Content

EndoC-βH1cells were seeded onto coated 24-well plates at a density of 250 × 10^3^ cells per well for 72 h before EDC treatment. Then, cells were exposed to the estrogenic agonists for 48 h. Prior to the insulin secretion experiment, cells were washed twice and preincubated for 3 h in a modified Krebs–Ringer medium containing 120 mM NaCl, 5.4 mM KCl, 1.2 mM KH_2_PO_4_, 1.2 mM MgSO_4_, 20 mM HEPES, 2.4 mM CaCl_2_ and 0.1% BSA, pH 7.4 at 37 °C. Cells were then incubated with Krebs–Ringer medium at 0 mM glucose for 40 min, followed by 20 mM glucose for 40 minutes. Supernatants were collected as basal and stimulated insulin secretions, respectively, centrifuged at 700 × g and 4 °C for 5 min, and immediately frozen for later analysis. For insulin content cells were lysed using TETG buffer supplemented with Complete™ Mini Anti-protease (Sigma-Aldrich, Saint Louis, MO, USA) (prepared following the manufacturer’s instructions). Lysate was centrifuged at 700× g and 4 °C for 5 min and supernatants were immediately frozen. Insulin secretion and content were measured by ELISA according to manufacturer’s instructions using the Human Insulin Kit (Mercodia, Uppsala, Sweden). Insulin released during each 40 minutes incubation was converted to an hourly rate and divided by the intracellular insulin content measured in the corresponding well after the secretion assay. Insulin content was expressed as a percentage of the paired vehicle control. Each of four independent experiments included three wells per condition; wells were averaged within each experiment before analysis. For simplicity, only significant differences due to ER agonist treatment between low or high-glucose and vehicle-treated cells are denoted.

### 5.5. RNA Extraction and RT-qPCR

EndoC-βH1 pancreatic cells were seeded in 24-well plates at a density of 200 × 103 cells/well and cultured for 72 hours. After agonist treatment, RNA was extracted using a commercial kit (RNeasy Micro kit, Qiagen, Hilden, Germany) according to the manufacturer’s instructions. RNA (1 μg) was reverse transcribed using the High Capacity cDNA Reverse Transcription kit (Applied Biosystems, Foster City, CA, USA). Quantitative PCR assays were performed using the CFX96 Real Time System (Bio-Rad Laboratories, Hercules, CA, USA). Amplification reactions were carried out in medium containing a 200 nM concentration of each primer, 1 μL of cDNA, and IQ SYBR Green Supermix (Bio-Rad Laboratories, Hercules, CA, USA). Primers were designed between exons to avoid genomic cross-reaction. Samples were subjected to the following conditions: 30 s at 95 °C, 45 cycles (5 s at 95 °C, 5 s at 60 °C, and 10 s at 72 °C) and a melting curve of 65–95 °C. The resulting values were analyzed with the CFX96 Real-Time System (Bio-Rad Laboratories, Hercules, CA, USA) and were expressed relative to the control values (2−ΔΔCT). The primers used for qPCR were purchased from Invitrogen (Thermo Fisher Scientific), and their sequences are: KCNJ11: Forward (5′–3′), CCATTGAGGTCCAAGTGACTATT; Reverse (5′–3′), ACGATGTTCTGCACGATGAG. KCNB1: Forward (5′–3′), GCTTTCGCTTTCTGCCTCGC; Reverse (5′–3′), GGCCGTCTTCTCACCTCCAT.KCNB2: Forward (5′–3′), GTCTGTGGGAGCTAGCTTTGT; Reverse (5′–3′), GCAGAGCCAGGGTCCTTAAA. KCNMA1: Forward (5′–3′), GACGCTCAAGTACCTGTGGA; Reverse (5′–3′), ACTAAGACAACCTTCGTCTTGCC. CACNA1H: Forward (5′–3′), CTGTTCGGGCAGAAGTGTTA; Reverse (5′–3′), TGTGTCCGTCCAACGAGTA. CACNA1A: Forward (5′–3′), ACAGTTGGGACGGAGTTTG; Reverse (5′–3′), CGACTTCAGGACGACTTGTAAA. CACNA1C: Forward (5′–3′), CCCTGCTTGTGCTGTTTGTC; Reverse (5′–3′), CCTCCTGGTTGTAGCAGGTC. CACNA1D: Forward (5′–3′), CGTTTGACTCCCTCATCGTAAT; Reverse (5′–3′), CAGTTGGGACAGGGACATTT. CACNA1E: Forward (5′–3′), CGAAGACTGGAGAAGACAGAAC; Reverse (5′–3′), AGAGCCCTTATGGAAGATGAAC. SCN9A: Forward (5′–3′), CTTTTCTTCGTGACCCGTGG; Reverse (5′– 3′), GCCCCTACAATTGTCTTCAGG. SCN8A: Forward (5′–3′), CTCCTGACTGGTCGAAGAATG; Reverse (5′–3′), AGCCATCTATGCAGAAACCTC. ACTB (β-actin): Forward (5′–3′), CACCATTGGCAATGAGCGGTTC; Reverse (5′–3′), AGGTCTTTGCGGATGTCCACGT. All measurements were performed in duplicate and normalized against the geometric mean of the housekeeping genes *Actb*.

### 5.6. Electrophysiological Recordings

EndoC-βH1 cells were plated on ECM–fibronectin pre-coated slip slides (10 mm) at a density of 200 × 10^3^ cells per slide. After at least 72 hours cells were exposed to agonists during 48 hours at 1 nM concentration. For the patch-clamp recordings of voltage-gated K^+^, Ca^2+^ and Na^+^ currents the standard whole-cell patch-clamp was used, as previously described (20, 30). Data were obtained using an Axopatch 200B amplifier (Axon Instruments Co., Union City, CA, USA). Patch pipettes were pulled from borosilicate capillaries (Sutter Instruments Co., Novato, CA, USA) using a flaming/brown micropipette puller P-97 (Sutter Instruments Co., Novato, CA, USA) and heat polished at the tip using an MF-830 microforge (Narishige, Japan). For recordings of voltage-gated K⁺ currents, the bath solution contained 138 mM NaCl, 5.6 mM KCl, 2.6 mM CaCl₂, 1.2 mM MgCl₂, 5 mM HEPES, and 5 mM glucose (pH: 7.4 with NaOH). Bath temperature was maintained at 32–35 °C in all recordings. Various pipette-filling solutions were used. For recordings of voltage-gated K⁺ currents, the pipette was filled with the following: 120 mM KCl, 1 mM MgCl₂, 1 mM CaCl₂, 3 mM MgATP, 10 mM EGTA, and 10 mM HEPES (pH: 7.2 with KOH). A similar internal solution was used for the Ca²⁺ current measurements, except that 120 mM KCl was replaced by 130 mM CsCl and pH was adjusted with CsOH (pH: 7.2). The bath solution used for Ca²⁺ current measurements contained 118 mM NaCl and 20 mM TEA-Cl, with the remaining components unchanged from the K⁺ recording solution, and 1 µM TTX was added to block voltage-gated Na⁺ currents. To measure Na⁺ currents, we used the same internal solution as for Ca²⁺ currents and a bath solution containing 118 mM NaCl, 20 mM TEA-Cl, 5.6 mM KCl, 2.35 mM CaCl₂, 1 mM MgCl₂, 5 mM HEPES, 5 mM glucose, and 250 µM cadmium chloride (CdCl₂) to block Ca²⁺ currents (pH: 7.4 with NaOH). The electrophysiological protocols for recording of K^+^, Ca^2+^ and Na^+^ currents are specified in the text and figure legends. For current density quantification, K^+^, Ca^2+^ and Na^+^ currents (in pA) were normalized to the cell capacitance (in pF).

After filling the pipette with the pipette solution, the pipette resistance was 3–5 MΩ. A tight seal (>1 GΩ) was established between the cell membrane and the tip of the pipette by gentle suction. The series resistance of the pipette usually increases to 6–15 MΩ after moving to the whole cell. Series resistance compensation was used (up to 40%) to keep the voltage error below 5 mV during current flow. Finally, data were filtered (2 kHz) and digitized (10 kHz) using a Digidata 1550B1 (Molecular Devices, San Jose, CA, USA).

For the perforated patch-clamp recordings, membrane potential was recorded in current-clamp mode at zero injected current. The pipette solution contained (mM): 128 K-gluconate, 10 KCl, 10 NaCl, 1 MgCl₂ and 10 HEPES (pH 7.35 with KOH), with 0.24–0.32 mg/mL amphotericin B. The bath contained (mM): 138 NaCl, 5.6 KCl, 2.6 CaCl₂, 1.2 MgCl₂, 5 HEPES and 0.5 glucose (pH 7.4). Cells were preincubated for 3 h without glucose and recorded at least 15 min after switching to 0.5 mM glucose. Recordings began after access resistance stabilized below 50 MΩ. Liquid junction potential was corrected, and recordings with unstable perforation or membrane rupture were excluded.

### 5.7. Statistical Analysis

The GraphPad Prism 8.0 software (GraphPad Software, Inc., San Diego, CA, USA) was used for all statistical analyses. Data are expressed as the mean ± SEM. Comparisons between two independent groups were performed using unpaired, two-tailed Student’s t-tests. Comparisons involving multiple groups were performed using ordinary or repeated-measures one-way ANOVA followed by Tukey’s or Dunnett’s multiple comparisons test, as appropriate. Two-way repeated-measures ANOVA followed by Dunnett’s multiple comparisons test was used to assess treatment effects across glucose concentrations or voltages, with Greenhouse–Geisser correction applied to current–voltage analyses. Statistical significance was set at p < 0.05. Sample sizes and the specific statistical tests used are indicated in the corresponding figure legends.

## Results

### Expression and location of ERα and ERβ

To study the expression and cellular localization of ERα and ERβ in EndoC-βH1 cells we performed immunocytochemistry using antibodies against both receptors. ERα which stained red, was uniformly distributed throughout the cell (Fig.1a), including the nucleus, which stained blue with Hoechst (Fig. 1b). ERβ, also stained red, was predominantly localized outside the nucleus, although weak staining was observed inside the nucleus, albeit to a lesser extent than in the cytoplasm (Fig. 1c and d). To further characterize the extranuclear localization of ERβ, we analyzed its colocalization with markers of intracellular organelles. ERβ was found to colocalize with KDEL, a marker of the endoplasmic reticulum (Fig. 1e), with a Manders colocalization coefficient (MCC) of 0.87 ± 0.04, GM130, a marker of the Golgi apparatus (MCC = 0.89 ± 0.07) (Fig. 1f), and cytochrome c (CytC), a mitochondrial marker (MCC = 0.74 ± 0.09) (Fig. 1g). These results suggest that ERβ is associated with multiple intracellular organelles, including the endoplasmic reticulum, the Golgi apparatus, and the mitochondria. This supports the idea that ERβ is predominantly located extranuclearly in EndoC-βH1 cells.

**Figure 1.**
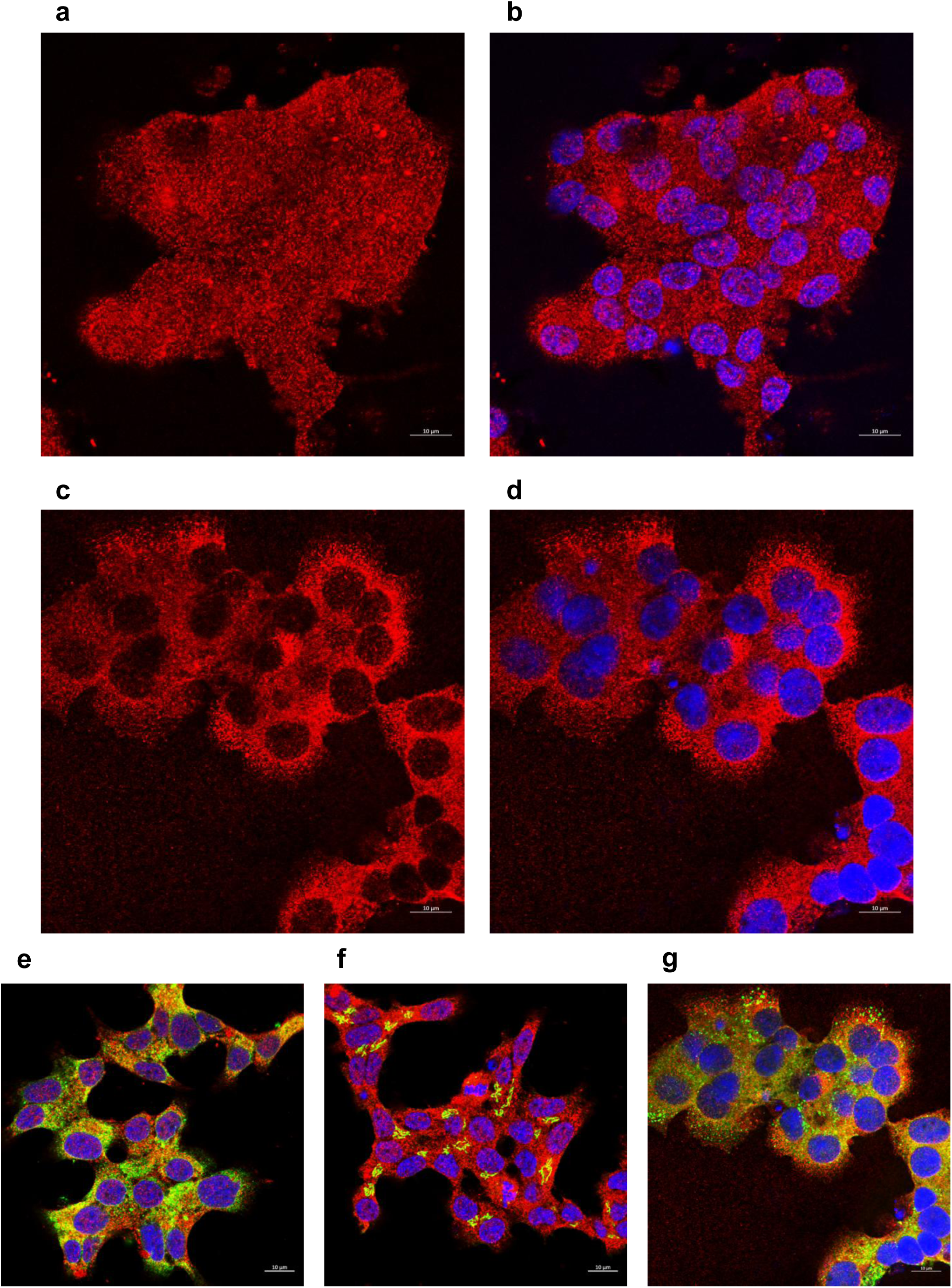
ERα is uniformly distributed throughout the cell while ERβ is preferentially located outside the nucleus in EndoC-βH1 cells. **(a)** Representative image showing the cellular distribution of ERα. **(b)** The nucleus, stained with Hoechst (in blue), is seen superimposed to ERα. **(c)** Representative image showing the cellular distribution of ERβ. **(d)** The nucleus, stained with Hoechst (in blue), is seen superimposed to ERβ. **(e)** Representative image showing the colocalization of ERβ with KDEL, an endoplasmic reticulum marker. **(f)** Representative image showing the colocalization of ERβ with GM130, a Golgi apparatus marker. **(g)** Representative image showing the colocalization of ERβ with cytochrome c (CytC), a mitochondrial marker. The calibration bar is 10 μm.

### Effect of E2 and DPN on insulin secretion

To study the effect of ERα and ERβ on GSIS, we treated EndoC-βH1 during 48 hours with PPT and DPN, as well as the hormone, 17β-estradiol (all at 1nM concentration). E2 and DPN treatments induced a decrease in the basal insulin secretion of 38% and 26%, respectively, compared to their matched controls. The GSIS in response to 20mM glucose from 0 mM was about 3-fold in control conditions but was unaffected by any treatment (Fig. 2a). The content of insulin was unaffected by any treatment (Fig. 2b). These results indicate that the decrease in basal insulin secretion is due to a regulation in the stimulus-secretion coupling rather than a decreased insulin production in human EndoC-βH1 cells.

**Figure 2.**
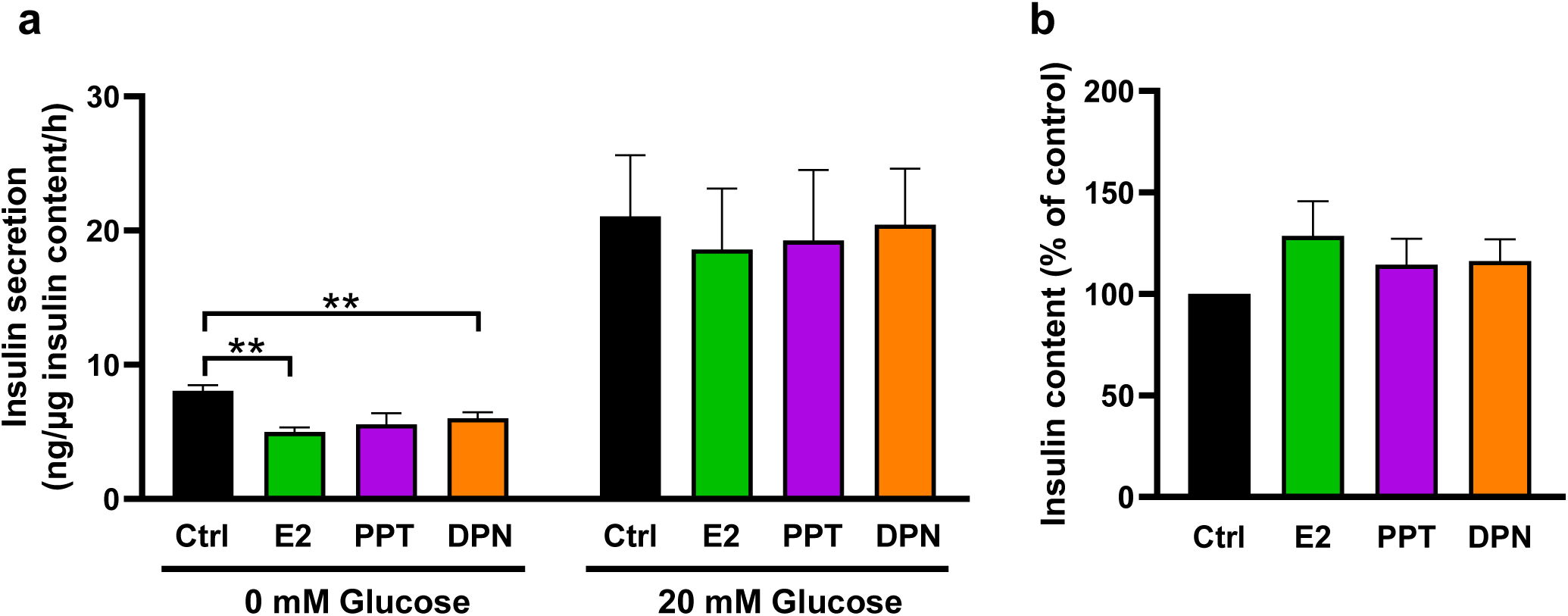
Estradiol and ligand of estrogen receptors DPN (ERβ) decrease the basal insulin secretion without altering insulin content in EndoC-βH1 cells. **(a)** Insulin secretion normalized to total insulin content under basal (0 mM) and stimulatory (20 mM) glucose conditions, expressed as ng insulin/µg insulin content/h. EndoC-βH1 cells were treated for 48 hours with vehicle control, 1 nM E2 (green), 1 nM PPT (violet), or 1 nM DPN (orange). **(b)** Insulin content expressed as percentage of the paired control condition for each independent experiment after 48 hours treatment with vehicle control (black), 1nM E2 (green), 1 nM PPT (violet) or 1 nM DPN (orange). Data are mean ± SEM; vs control (RM One-Way ANOVA; Dunnett’s multiple comparisons); n = 4 independent experiments. **p<0.01.

### Effects of PPT and DPN on electrical activity

To study the effect of ERs activation on the stimulus-secretion coupling, we sought to investigate the effect of PPT and DPN on the electrical activity recording the membrane potential using the perforated patch-clamp technique (see details in Material and methods section). Following 3 hours of preincubation in glucose-free recording solution and at least 15 min of equilibration at 0.5 mM glucose, control EndoC-βH1 cells fired action potentials at a mean frequency of 0.81 ± 0.07 Hz (Fig. 3a, d). These recording conditions were used to assess the effects of PPT and DPN on electrical activity (Fig. 3a–e). Treatment with the ERα agonist, 1 nM PPT for 48 hours, increased the frequency of action potential firing compared with vehicle-treated cells (1.07 ± 0.04 Hz) (Fig. 3a, b, d). The increased firing was induced, even to a greater extent, by the ERβ agonist, 1 nM DPN (1.38 ± 0.05 Hz) (Fig. 3a, c, d). The resting membrane potential (RMP) of cells treated during 48 hours with 1 nM PPT or 1 nM DPN was more depolarized (−54.7 ± 1.1 mV and −55.5 ± 1.8 mV, respectively), than vehicle-treated cells (−64.1 ± 0.7 mV) (Fig. 3a,b,c,e). When we analyzed the action potential shape, we observed that both PPT and DPN reduced their amplitude (Fig. 4a,b), however the half-width was enlarged (Fig. 4a,c). These changes were associated with an increased area under the curve of the action potential (Fig. 4a,d), indicating that the electrical charge transferred through the plasma membrane during an action potential is increased. In pancreatic β-cells, as in other excitable cells, action potential firing results from a complex interplay between voltage-dependent and voltage-independent ionic conductance. We sought then to investigate the effects of specific activation of ERs on ion channel expression and currents.

**Figure 3.**
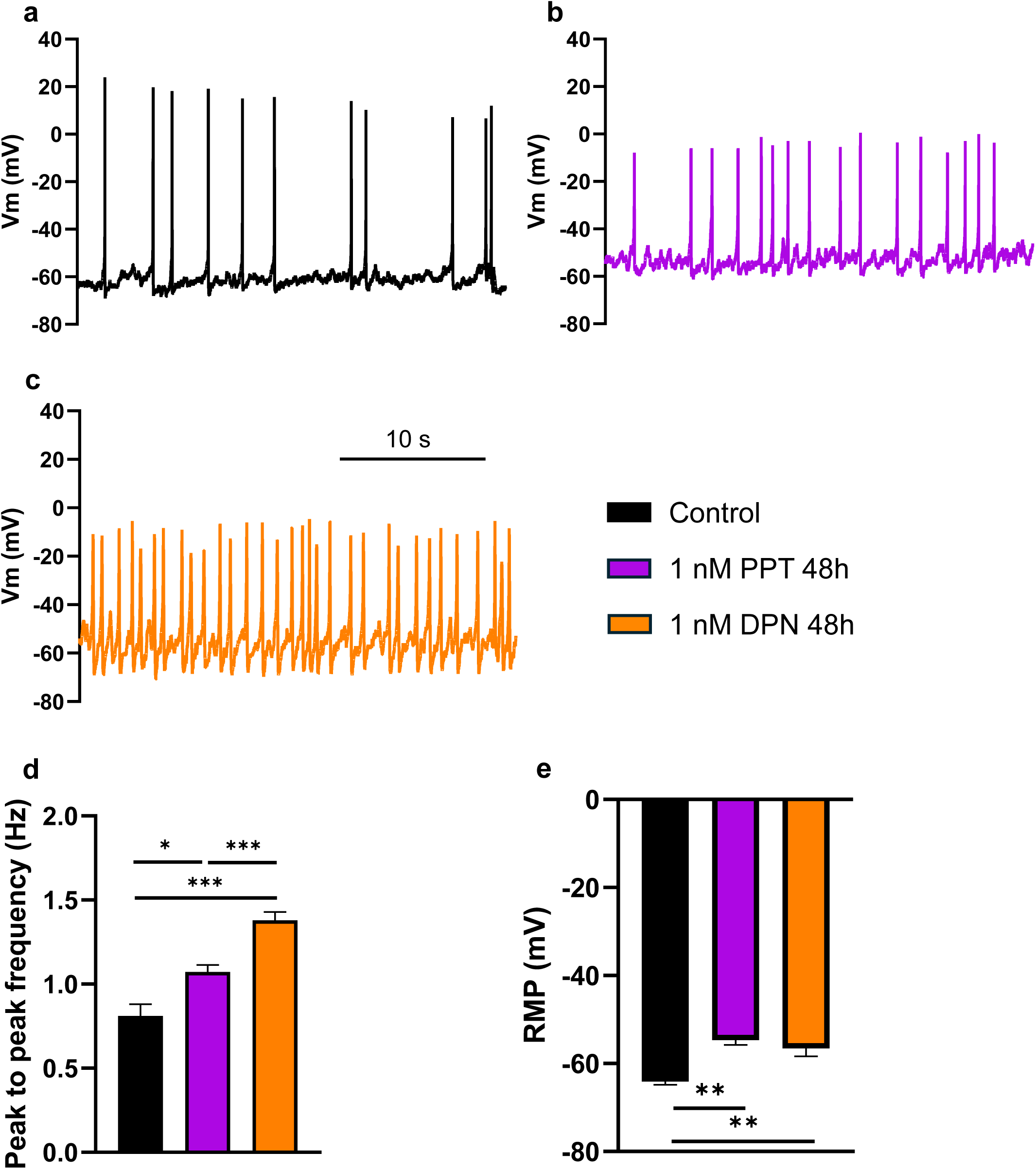
PPT and DPN modify electrical activity in EndoC-βH1 cells. **(a-c)** Representative recordings of action potential firing of EndoC-βH1 cells treated for 48 hours with control (black), 1 nM PPT (violet) or 1 nM DPN (orange) at 0.5 mM glucose. Time bar: 10 s. **(d)** Peak to peak action potential frequency of control (black), PPT (violet) and DPN (orange). **(e)** Average resting membrane potential of control (black), PPT (violet) and DPN (orange). Data are mean ± SEM; *p < 0.05, **p < 0.01, ***p < 0.001 (One-Way ANOVA; Tukey’s multiple comparisons). Control n = 5, PPT n = 4 and DPN n = 5 cells. pooled from 5 independent experiments.

**Figure 4.**
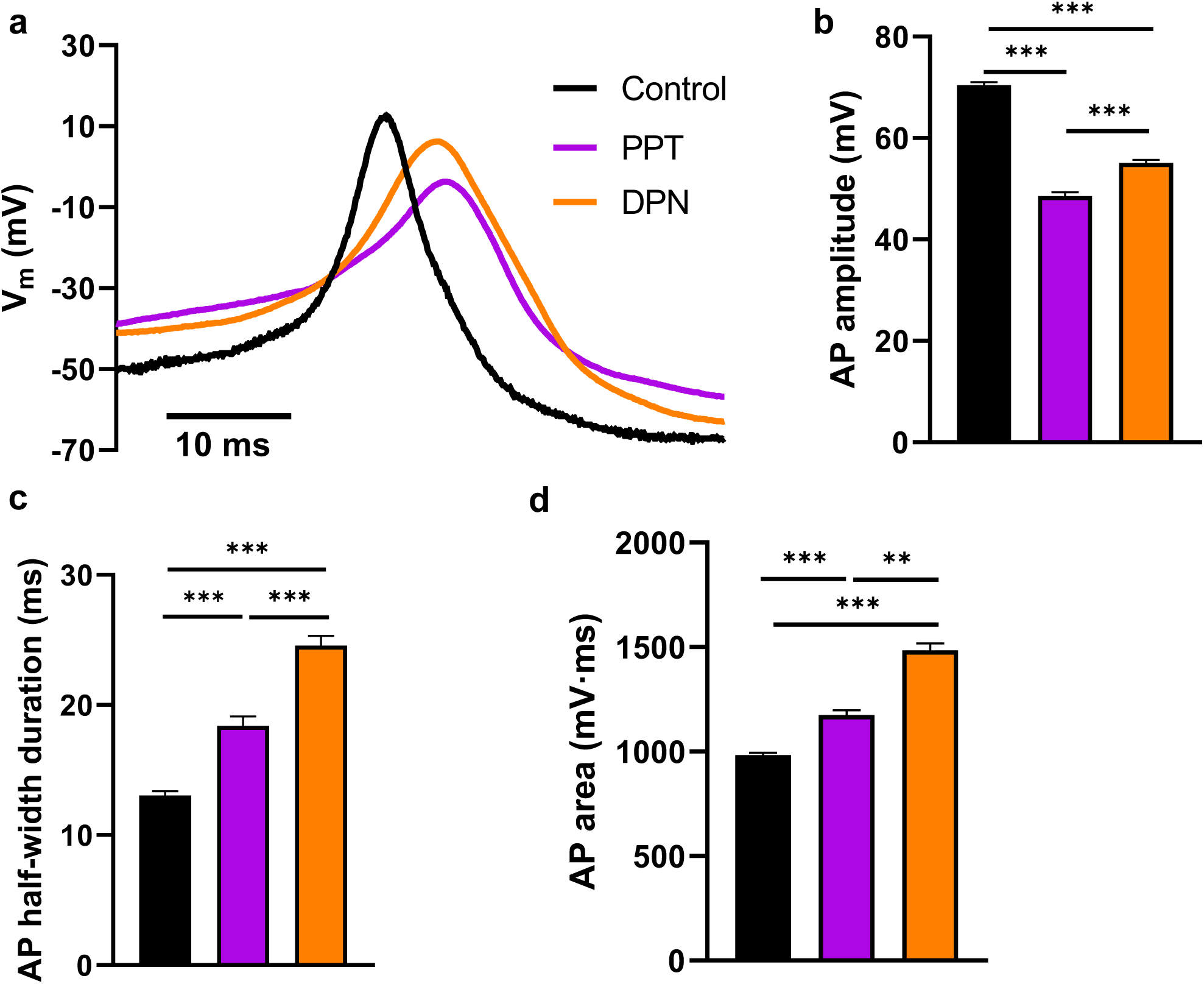
PPT and DPN modify the amplitude and duration of action potentials in EndoC-βH1 cells. **(a)** Representative recording of a single action potential in EndoC-βH1 cells treated for 48 hours with control (black), 1 nM PPT or DPN at 0.5 mM glucose. **(b)** Action potential amplitude of control (black), PPT (violet) and DPN (orange). **(c)** Action potential half-width duration of control (black), PPT (violet) and DPN (orange). **(d)** Action potential area under the curve (effective charge transfer) in control (black), PPT (violet) and DPN (orange). Data are mean ± SEM; **p < 0.01, ***p < 0.001 (One-Way ANOVA; Tukey’s multiple comparisons). Control n = 5 cells,

### No effect of PPT and DPN on the gene expression of channel subunits

To study the effects of ERα or ERβ activation on gene expression in EndoC-βH1 cells, we treated these cells for 48 hours with 1 nM of E2, PPT or DPN, and performed quantitative RT-PCR to measure the expression level of the main ion channel genes in pancreatic β-cells.

EndoC-βH1 cells are equipped with K^+^ channels, including the ATP-sensitive K^+^ channel Kir6.2 (which is encoded by *KCNJ11*), the voltage-gated K^+^ channels delayed rectifiers Kv2.1 (*KCNB1*) and Kv2.2 (*KCNB2*), and the large-conductance Ca^2+^-activated K^+^ channel KCa1.1/BK channel (*KCNMA1*). These channels are involved in establishing the resting membrane potential, the repolarization of the action potential and controlling firing rate, respectively. Treatment with E2, PPT, or DPN for 48 hours at 1nM did not affect the expression of any of these K⁺ ion channel subunits (Fig. 5a–d).

**Figure 5.**
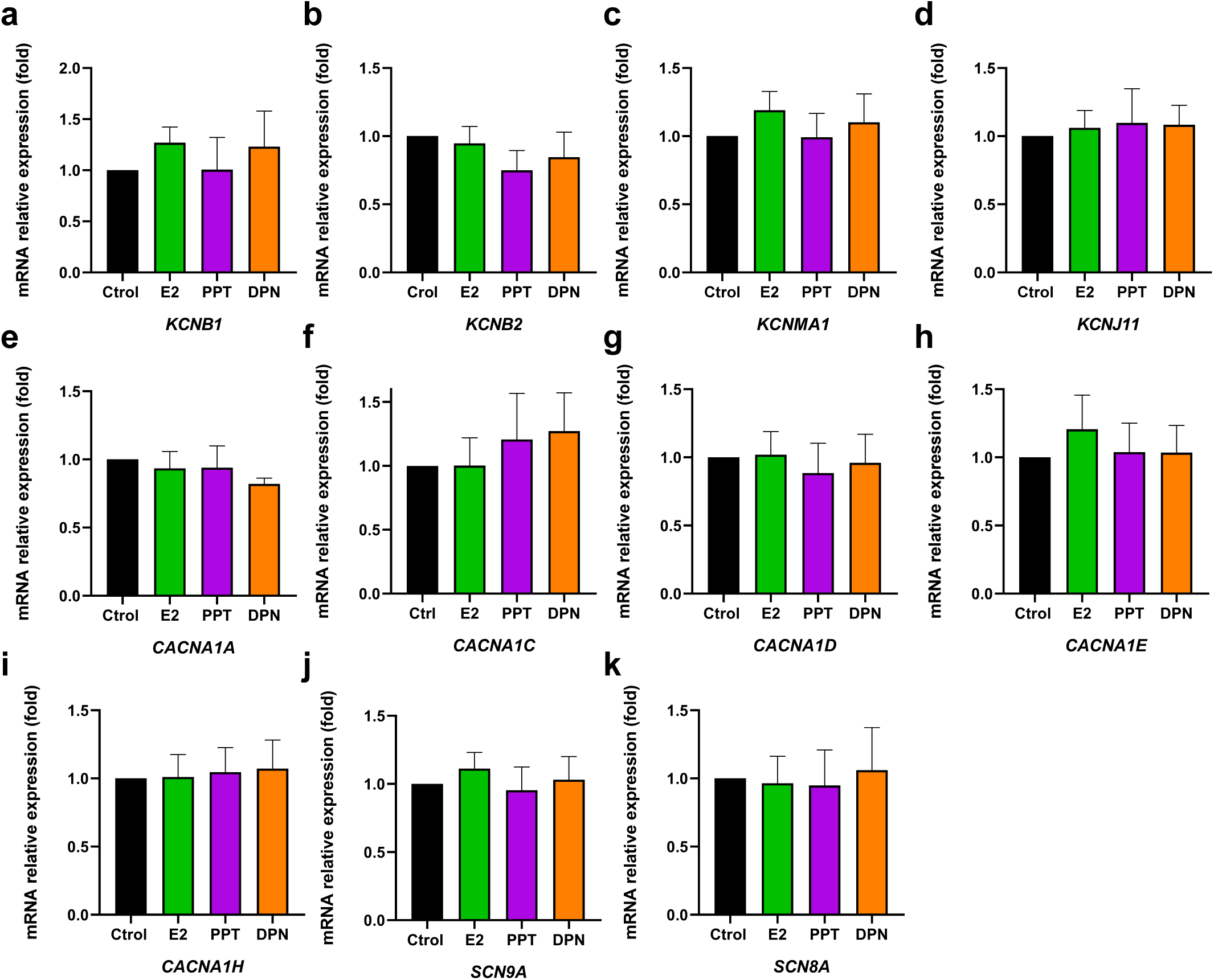
Lack of effect of estrogen receptor ligands on ion channel expression in EndoC-βH1 cells. The expression of the following ion channel subunits, measured by mRNA expression levels, were unaffected by E2, PPT or DPN treatment during 48 hours at 1 nM. K^+^ channels: **(a)** Kv2.1 (*KCNB1*), **(b)** Kv2.2 (*KCNB2*), **(c)** KCa1.1/BK channel (*KCNMA1*) and **(d)** KATP (*KCNJ11*). Ca^2+^ channels: **(e)** P/Q-type (*CACNA1A*), **(f-g)** L-types (*CACNA1C* and *CACNA1D*), **(h)** R-type (*CACNA1E*) and **(i)** T-type (*CACNA1H*). Na^+^ channels: **(j)** Nav1.7 (*SCN9A*) and **(k)** Nav1.6 (*SCN8A*). Data are mean ± SEM; vs control (RM One-Way ANOVA; Dunnett’s multiple

EndoC-βH1 cells express low-voltage-activated T-type and high-voltage activated Ca^2+^ channels. T-type channels Cav3.2 (encoded by *CACNA1H*) are important in generating pacemaking electrical activity. High-voltage activated Ca^2+^ channels are composed of P/Q-type Cav2.1 (*CACNA1A*), L-type Cav1.2 (*CACNA1C*) and Cav1.3 (*CACNA1D*), and R-type Cav2.3 (*CACNA1E*), participate in insulin secretion. The expression of genes encoding for these channels was unaffected by E2, PPT and DPN treatment (Fig. 5e-i).

EndoC-βH1 cells have a large tetrodotoxin-sensitive Na^+^ current (24). The main Na^+^ channel α-subunits in these cells are the voltage-gated Nav1.7 (encoded by *SCN9A*) and the voltage-gated Nav1.6 (*SCN8A*) (24). The expression of both Na^+^ channel genes was unaffected (Fig. 5j,k).

These results point to the fact that the alterations in electrical activity induced by ER agonists was not due to changes in the expression of genes codifying for ion channels in EndoC-βH1 cells.

The electrophysiological changes observed after 48 hours treatment with ER agonists (Figs. 3–4 and 6–8) were not accompanied by statistically significant differences in the abundance of the selected ion-channel transcripts in EndoC-βH1 cells (Fig. 5a–k).

**Figure 6.**
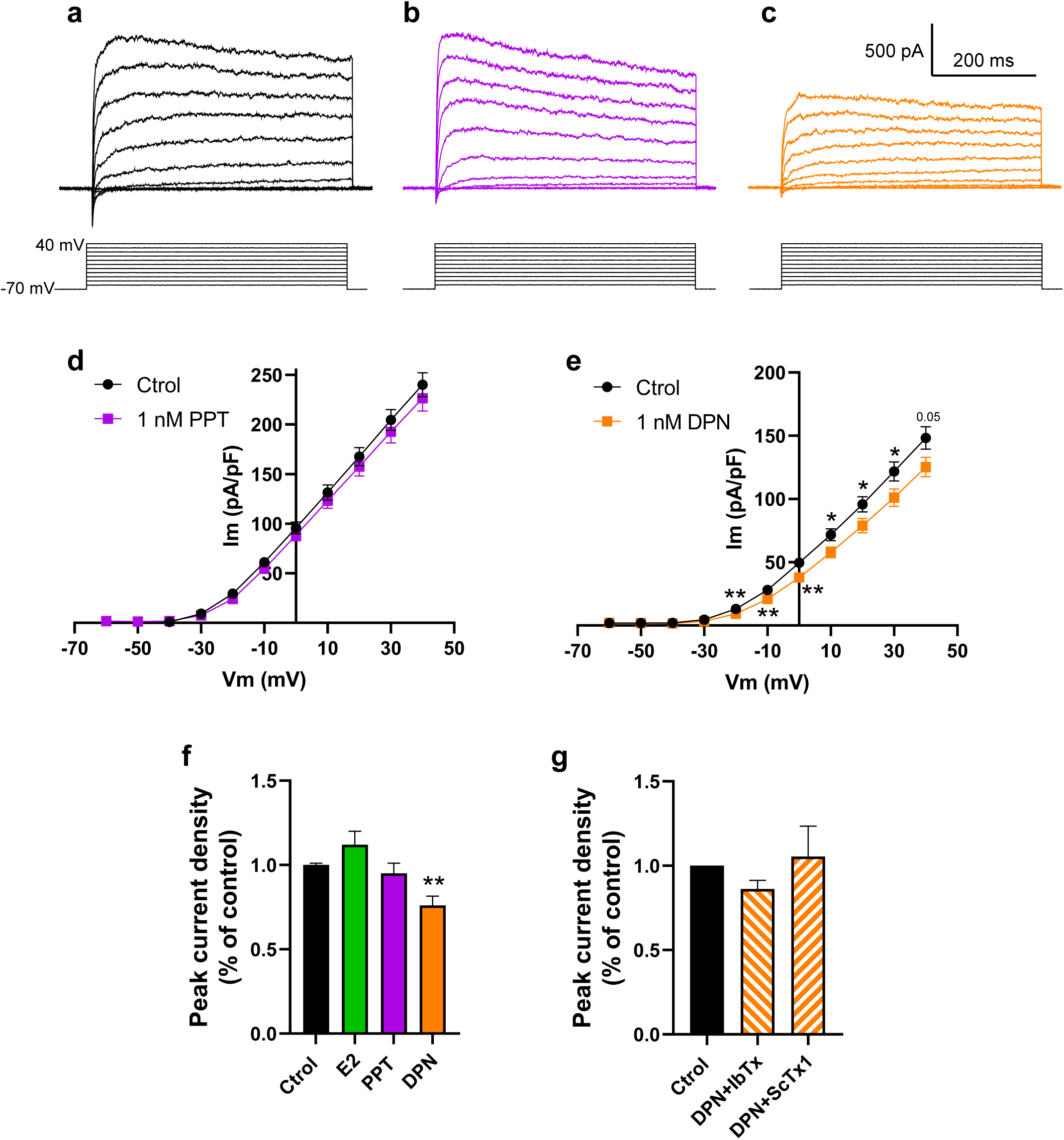
DPN inhibits K^+^ currents in EndoC-βH1 cells, but E2, and PPT are without effect. **(a–c)** Representative whole-cell K⁺ currents evoked by 10-mV depolarizing steps from −70 to +40 mV (500-ms pulses; voltage protocol shown beneath traces) in vehicle (black), 1 nM PPT (violet) and 1 nM DPN (orange) for 48 hours. Scale bars: 500 pA, 200 ms. **(d-e)** I–V relationships (pA/pF) for 1 nM PPT (violet) and 1 nM DPN (orange), vs their matched controls (black). Data are mean ± SEM; *p < 0.05, **p < 0.01 vs control at the indicated voltages (Student’s t-test). PPT panel: control n = 44 cells, PPT n = 45 cells, pooled from 4 independent experiments. DPN panel: control n = 32 cells, DPN n = 30 cells, pooled from 4 independent experiments. **(f)** Peak K^+^ current density at 0 mV expressed as % of control for all ligands (bars: mean ± SEM). E2 bar: control n = 32 cells, E2 n = 30 cells, pooled from 4 independent experiments. PPT bar: control n = 30 cells, PPT n = 30 cells, pooled from 4 independent bar. DPN bar: control n = 44 cells, DPN n = 45 cells, pooled from 4 independent experiments. **(g)** Peak K^+^ current density at 0 mV (pA/pF) for 1 nM DPN 48 hours + 100 nM IbTX or 100 nM ScTX1 versus its matched DMSO control (bars: mean ± SEM). Data are mean ± SEM; (Student’s t test). IbTX bar: control n = 22 cells, DPN n = 20 cells, pooled from 6 independent experiments. ScTX1 bar: control n = 28 cells, DPN n = 25 cells, pooled from 6 independent experiments.

### Actions on K^+^ channel-mediated currents

Next, we investigated whether ER activation modified ion channel currents. Using the patch-clamp technique in the whole-cell configuration we recorded voltage-dependent K^+^ channels in cells with 1 nM E2, PPT, or DPN for 48 hours. Figure 6 shows the recordings of the K^+^ currents in response to depolarizing voltage pulses ranging from −60 mV to +40 mV from a holding potential of −70 mV in cells treated with vehicle, 1 nM PPT or 1 nM DPN (Fig. 6a-c). Measurements of the whole-cell K^+^ currents revealed that PPT and E2 did not affect peak current density (Fig. 6d,f). In contrast, the currents were significantly smaller in DPN-treated cells than in vehicle-treated cells (Fig. 6e,f). The agonist of GPER, G1 (100 nM, 48h) had no effect on K^+^ currents (unpublished result). The effect of DPN supports the idea that ERβ is involved in the regulation of K^+^ currents as previously shown in mice (20, 30).

To determine the effect of DPN treatment (1 nM during 48 hours) on specific K⁺ channel components, we measured the macroscopic K⁺ currents in the absence or presence of selective blockers of the K⁺ currents identified in EndoC-βH1 cells. Macroscopic K⁺ currents were recorded before and after the application of iberiotoxin (IbTX, 100 nM), which inhibits BK channels, allowing isolation of the Kv current component. We also applied stromatoxin-1 (ScTX1, 100 nM), which inhibits Kv-family channels, allowing isolation of the BK current component (Fig. 6g). Treatment with DPN did not significantly affect the Kv or BK current components measured in the presence of the toxins. These results suggest that the functional changes induced by DPN are unlikely to be due to a significant modulation of these subtypes of K⁺ channels.

### Actions on Ca^2+^ channel-mediated currents

To test whether PPT and DPN could regulate voltage-dependent Ca^2+^ entry in EndoC-βH1 cells, we incubated the cells with agonists for 48 hours. Then, we studied the effects of specific ER activation on Ca^2+^ currents using the patch-clamp technique in the whole-cell mode.

Figure 7 shows representative recordings of the Ca^2+^ currents in response to depolarizing voltage pulses ranging from −60 mV to +60 mV from a holding potential of −70 mV in EndoC-βH1 cells treated with vehicle, 1 nM PPT or 1 nM DPN (Fig. 7a-c). As can be observed in Fig. 7d, incubation with 1 nM PPT increased the voltage-gated Ca^2+^ currents at several holding potentials including those of the action potential shape. In contrast, incubation with 1 nM DPN had no effect on voltage-gated Ca^2+^ currents. We also measured the apparent voltage dependence of Ca^2+^ current activation that was estimated by fitting individual peak current density–voltage relationships with a modified Boltzmann equation. The V_1/2_ of activation values were: Control: −22.39 ± 0.83 mV; PPT: −19.17 ± 0.88 mV and DPN: −20.18 ± 0.90 mV. PPT 1 nM for 48 hours significantly shifted the apparent V_1/2_ of activation toward more depolarized potentials (Fig. 7e). DPN showed a non-significant trend in the same direction (Fig. 7e). Taken together, these results demonstrate that selective activation of ERα with PPT, but not ERβ with DPN, enhances voltage-gated Ca²⁺ currents and shifts the apparent activation V₁/₂ towards more depolarized potentials in human EndoC-βH1 cells. The absence of an effect of DPN on Ca²⁺ currents suggests that ERβ activation does not promote insulin secretion through modulation of voltage-gated Ca²⁺ channel activity.

**Figure 7.**
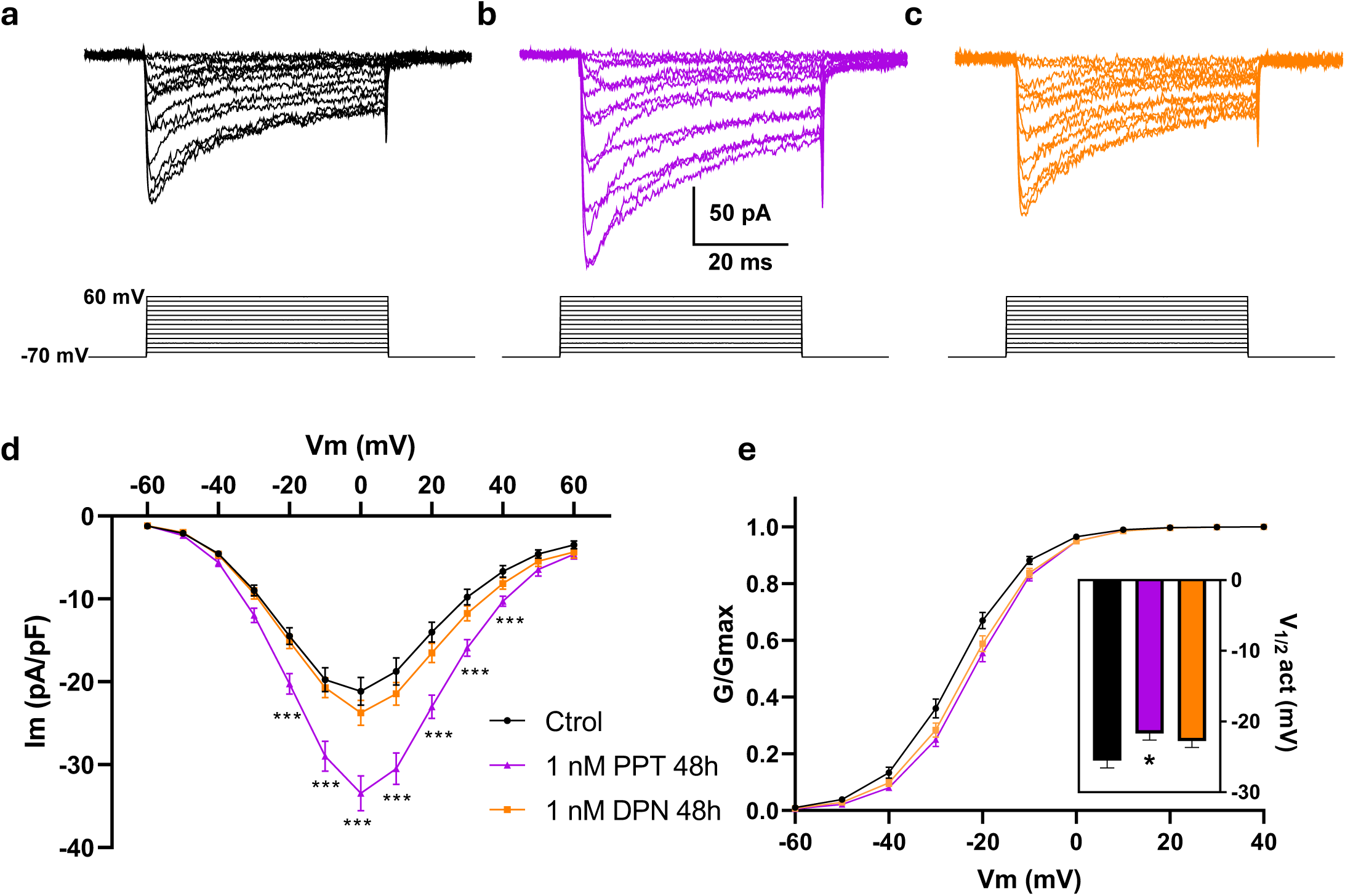
PPT increases Ca^2^⁺ currents in EndoC-βH1 cells, but DPN is without effect. **(a-c)** Representative whole-cell Ca^2^⁺ currents evoked by 10-mV depolarizing steps from −70 to +60 mV (50-ms pulses; voltage protocol shown below current traces) in vehicle (black), 1 nM PPT for 48 h (violet) and 1 nM DPN for 48h (orange). Scale bars: 50 pA, 10 ms. **(d)** Current–voltage (I–V) relationships (current density, pA/pF) comparing PPT and DPN vs control. **(e)** Voltage dependence of Ca²⁺ current activation, represented as normalized conductance G/G_max_ against Vm. Data are mean ± SEM; *p < 0.05, **p < 0.01, ***p < 0.001, ****p < 0.0001 vs control at the indicated voltages. For I–V relationships in panel **d**, statistical comparisons were performed using repeated-measures two-way ANOVA with Greenhouse–Geisser correction followed by Dunnett’s multiple comparisons test versus control at the indicated voltages. For V_1/2_ values in panel **e**, statistical comparisons were performed using ordinary one-way ANOVA followed by Dunnett’s multiple comparisons test versus control. Control n = 21 cells, PPT n = 17 cells, DPN n = 18, pooled from 4 independent experiments.

### Actions on Na^+^ channel-mediated currents

The most abundant voltage-gated Na⁺ channel subtypes in EndoC-βH1 cells and human β-cells have been identified as Nav1.7 and Nav1.6 (24). The next experiment sought to investigate whether ER activation modulates Na⁺ channel function. Figure 8 shows representative recordings of the Na^+^ currents in response to depolarizing voltage pulses ranging from −60 mV to +60 mV from a holding potential of −70 mV in EndoC-βH1 cells treated with vehicle, 1 nM PPT or 1 nM DPN (Fig. 8a-c). Using this protocol, we observed a decrease in Na^+^ currents in cells treated with 1 nM DPN for 48 hours at −10 mV (Fig. 8d). However, this protocol does not account for the fraction of Na^+^ channels that are inactive at the holding potential of −70 mV. Therefore, we used an alternative protocol which is commonly employed in β-cells, to include the fraction of Na^+^ channels that inactivate at the holding potential of −70 mV (31). This protocol consists of applying a preceding 50 ms hyperpolarizing pulse (Vm = −150 mV). Then, we obtained a decrease in Na^+^ currents between −10 and +20 mV compared to matched controls (Fig. 8e). Unlike previous electrophysiological reports on EndoC-βH1 Na^+^ currents (24), we did not observe significant differences in the currents elicited from −70 mV and those elicited with the hyperpolarizing prepulse. In contrast to activation of ERβ with DPN, activation of ERα with PPT had no effect on Na^+^ currents in any of the protocols.

**Figure 8.**
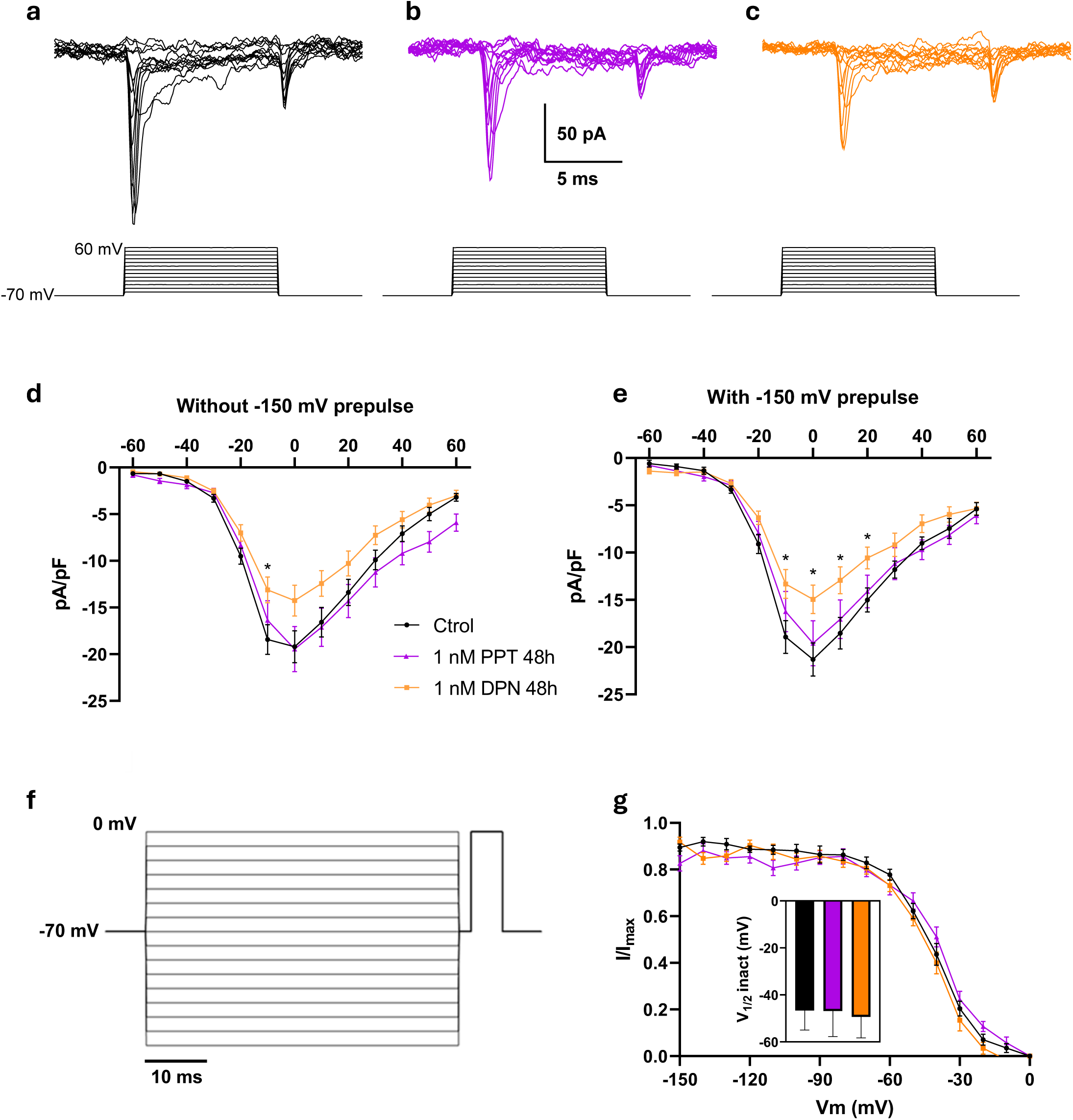
DPN decreases Na⁺ currents in EndoC-βH1 cells, but PPT is without effect. **(a-c)** Representative whole-cell Na⁺ currents evoked by 10-mV depolarizing steps from −70 to +60 mV (10-ms pulses; voltage protocol shown below current traces) in control (black), 1 nM DPN for 48 h (orange) and 1 nM PPT for 48 hours (violet). Scale bars: 50 pA, 5 ms. **(d)** Current–voltage (I–V) relationships (current density, pA/pF) comparing PPT and DPN vs control from the resting membrane potential. **(e)** Current– voltage (I–V) relationships (current density, pA/pF) comparing DPN and PPT vs control from the hyperpolarizing −150 mV voltage pulse. **(f)** Steady-state inactivation voltage protocol showing prepulses from −150 to 0 mV followed by a depolarizing pulse up to 0 mV. **(g)** Steady-state inactivation curves of I_Na_, represented as normalized Na⁺ channel availability (I/I_max_) after each conditioning prepulse. Data are mean ± SEM; *p < 0.05 vs control at the indicated voltages. For I–V relationships in panels **d**–**e**, statistical comparisons were performed using RM two-way ANOVA with Greenhouse–Geisser correction followed by Dunnett’s multiple comparisons test vs control at the indicated voltages. For the steady-state inactivation analysis in panel **g**, V_1/2_ values were obtained from individual Boltzmann fits and compared between groups using ordinary one-way ANOVA followed by Dunnett’s multiple comparisons test versus control. Control n = 13 cells, DPN n = 15, PPT n = 16 cells, pooled from 4 independent experiments.

Finally, to characterize the half-voltage of activation and the steady state of inactivation of Na^+^ channels, we applied a protocol consisting of a series of 50 ms, 10 mV voltage steps from −150 mV up to 0 mV, followed immediately by a 5 ms depolarizing pulse up to 0 mV, in order to measure the steady-state inactivation at various voltages (Fig. 8f). No significant shifts in the apparent voltage V_1/2_ of activation were observed either from −70 mV or after the −150 mV hyperpolarizing prepulse. Activation V₁/₂ values were obtained from fits to individual peak current–voltage relationships corresponding to Fig. 8d,e. From a holding potential of −70 mV, activation V₁/₂ values were: Control: −17.95 ± 0.94 mV; PPT: −16.16 ± 1.28 mV; and DPN: −17.83 ± 1.30 mV. After the −150 mV prepulse, activation V₁/₂ values were: Control: −16.87 ± 0.98 mV; PPT: −16.42 ± 1.02 mV; and DPN: −17.61 ± 1.55 mV. Regarding steady-state inactivation, V₁/₂ values were obtained from individual Boltzmann fits and compared by one-way ANOVA followed by Dunnett’s multiple comparisons test. No statistically significant differences were detected relative to control (Fig. 8g). Inactivation V₁/₂ values were: Control: −46.62 ± 2.31 mV; PPT: −46.87 ± 2.71 mV; and DPN: −49.37 ± 2.30 mV (Fig. 8g, inset).

Taken together, these results indicate that 1 nM DPN for 48 hours decreases Na^+^ current density within the voltage range at which Na^+^ currents reach their maximum, whether from a holding potential of −70 mV or following a hyperpolarizing −150 mV prepulse. However, this reduction was not accompanied by significant changes in the apparent voltage dependence of Na^+^ current activation or Na^+^ steady-state inactivation, as indicated by the lack of significant differences in V_1/2_ values between groups. This reduction in Na⁺ current density could decrease membrane excitability, representing a potential mechanism by which ERβ activation affects electrical activity and insulin secretion.

## Discussion

Our research indicates that both estrogen receptors (ERα and ERβ) are expressed in human insulin-secreting EndoC-βH1 cells. They are localized in both the nucleus and outside of it in the absence of added estrogenic compounds. Using the agonists PPT for ERα and DPN for ERβ, we found that activation of these estrogen receptors regulates ion channels and electrical activity in a manner that is distinct from changes in gene expression at the exposure time studied (48 hours). In addition, E2 and the specific ERβ agonist, DPN, both reduced the basal insulin secretion.

The limited availability of human islets and donor variability has hindered the study of human β-cell biology. Recent studies have shown that the human β-cell line, EndoC-βH1, resembles native human β-cells and therefore serves as a good model for investigating its physiology, including the study of stimulus-secretion coupling (24–26). The results presented in this study indicate that E2 and DPN decrease basal insulin secretion in EndoC-βH1 cells, as opposed to the effect described in mouse islets of Langerhans (3). Importantly, this difference highlights the value of studying human β-cell models directly, as the effects of ER activation observed in rodent models cannot necessarily be extrapolated to human β-cell physiology. This reduced effect on GSIS in EndoC-βH1 cells should be interpreted with caution, as this human cell line displays some features of an immature β-cell phenotype, including relatively high basal insulin secretion and a limited glucose-response window (32). The different experimental conditions used to assess insulin secretion, and EndoC-βH1 electrophysiology should be considered when interpreting our findings. In the insulin secretion experiments, stimulation with 20 mM glucose may have elicited a near-maximal secretory response, potentially limiting the ability to detect additional effects of PPT or DPN. By contrast, electrical activity was recorded at 0.5 mM glucose, under which EndoC-βH1 cells exhibited a relatively high basal action potential firing frequency. This suggests that estrogen receptor activation can further modulate β-cell excitability even under conditions of low glucose availability, although such changes do not necessarily translate into increased insulin secretion. Furthermore, voltage-gated ion currents were recorded at 5 mM glucose, providing an intermediate, physiologically relevant metabolic context in which the effects of ERα and ERβ activation on ion channel function could be characterized independently of the secretory response to high glucose. Overall, these differences highlight the importance of glucose concentration and the pre-existing functional state of EndoC-βH1 cells when comparing electrophysiological responses with insulin secretion.

Pancreatic β-cells couple stimuli with insulin secretion through changes in membrane potential and action potential firing, which regulate Ca^2+^ entry and exocytosis. Activation of ERα and ERβ affects stimulus-secretion coupling by regulating ion channel function rather than through gene expression. Gene expression of ion channels does not appear to play a significant role because no changes were observed in the expression of major genes encoding for Na^+^, Ca^2+^ and K^+^ channels. Nonetheless, we cannot discard changes in the expression of regulatory subunits that we did not study or changes in gene expression at shorter times of exposure. In any case, the change in the shape of the action potential and its consequences in the bursting pattern of electrical activity is most likely due to the regulation of ion channel function which is affected by both agonists, PPT and DPN.

The activation of ERα with PPT resulted in an increase in whole-cell Ca²⁺ current, while ERβ activation with DPN led to a reduction in K⁺ and Na⁺ currents. Despite these divergent effects, both agonists caused a depolarization of the resting membrane potential, an increase in action potential firing, and a prolongation of the action potential. These findings imply that ERα and ERβ may modulate β-cell stimulus-secretion coupling through receptor-specific remodeling of different ion-channel populations, rather than through a common mechanism. The convergence of these distinct effects on increased electrical activity may represent an important feature of estrogenic regulation of human β-cell excitability.

The increase in Ca^2+^ current produced by PPT suggests that ERα can directly modulate voltage-dependent Ca^2+^ entry in human β-cells. However, increased macroscopic Ca^2+^ current under voltage-clamp conditions does not necessarily imply proportionally greater Ca^2+^ influx during spontaneous action potentials, because PPT also depolarized the resting membrane potential, which may reduce channel availability and may prevent a proportional Ca²⁺ influx during spontaneous action potentials. Since the specific channel subtype remains unidentified, the exact contribution of this enhanced Ca²⁺ conductance to insulin secretion needs further research.

DPN produced a different electrophysiological phenotype, characterized by reductions in total outward K^+^ currents and Na^+^ inward currents. The decrease in K^+^ conductance is consistent with the increased action potential duration observed with DPN, since delayed-rectifier K^+^ channels contribute to repolarization in EndoC-βH1 cells.

Reduced outward K⁺ current may contribute to action-potential prolongation, but its consequences for Ca²⁺ entry and insulin secretion remain unresolved. No statistically significant increase in secretion at 20 mM glucose was detected. However, pharmacological dissection did not identify a specific K^+^ channel subtype responsible for the DPN effect. Therefore, the reduction in K^+^ current should be considered a plausible contributor, rather than a demonstrated cause, of the DPN-induced electrical phenotype.

Protecting the functional mass of pancreatic β-cells is a therapeutic challenge in patients with diabetes mellitus. Activation of ERs by estrogens improves insulin synthesis, GSIS and protects β-cells from apoptosis (3, 4, 10). In addition, women assigned with menopausal estrogen therapy showed improved glucose homeostasis (33–36). The challenge of using estrogens for diabetes therapy has been the risk of hormone-dependent cancers when ERs are activated. A novel strategy to retain the metabolic benefits of estrogen while reducing systemic adverse effects is the use of GLP-1–E2 conjugates. These compounds have demonstrated positive effects on body weight, glucose metabolism and insulin sensitivity in preclinical models, while reducing the uterotrophic effects associated with unconjugated E2 (37–40). GLP-1-E2 conjugates have recently demonstrated efficacy in delaying diabetes onset and preserving β-cell mass in female NOD mice (41). These conjugates are designed to target specific tissues that express GLP-1 receptors, avoiding E2 alone to enter into the circulation and thereby, reducing the risk of hormone-dependent cancer. However, this remains a concern for their use. GLP-1 conjugates incorporating ERβ-selective agonists may represent a promising alternative to GLP-1–E2 conjugates for diabetes treatment, potentially retaining the beneficial effects of ERβ activation on pancreatic β-cell function while minimizing the risks associated with systemic estrogen exposure.

The role of estrogen receptor subtypes in cancer research has been a subject of significant interest, particularly in the context of estrogen-dependent cancers. Research indicates that ERβ triggers apoptosis in breast cancer cells, unlike ERα, which stimulates cell division (42, 43). In our study, activation of ERs did not enhance glucose-stimulated insulin secretion at 20 mM glucose but rather reduced basal insulin secretion under glucose-free conditions (0 mM glucose). These findings highlight the complex and context-dependent effects of estrogen receptor activation on human β-cell function, which may differ from the effects on β-cell survival, β-cell proliferation, and glucose homeostasis reported in animal models of type 1 and type 2 diabetes (4, 44).

In summary, our results provide evidence that both ER subtypes can contribute to the regulation of stimulus-secretion coupling in human β-cells and highlight the potential of receptor-selective estrogenic signaling as a strategy to modulate β-cell function.

## Statements

### Data availability statement

The raw data supporting the conclusions of this article will be made available by the authors, without undue reservation.

### Author contributions

AN and JM-P conceived the study. RS-N, TE, CQ, MS-M and JM-P collected and analyzed the data. RS-N, TE, CQ, MS-M, AN y JM-P interpreted the data. RS-N, AN and JM-P drafted the manuscript. AN and JM-P supervised the study. RSN, AN and JM-P critically revised the manuscript for important intellectual content. JM-P and AN acquired the funding. All authors reviewed and approved the final version of the manuscript and gave consent for publication. AN and JM-P are responsible for the integrity of the work as a whole.

## Acknowledgements

The authors thank Maria Luisa Navarro, Salomé Ramon, and Beatriz Bonmati Botella for their excellent technical assistance.

## References

1. Duncan BB, Magliano DJ, Boyko EJ. IDF Diabetes Atlas 11th edition 2025: global prevalence and projections for 2050. Nephrol Dial Transplant. 2025;41(1):7–9.

2. Nadal A, Rovira JM, Laribi O, Leon-quinto T, Andreu E, Ripoll C, et al. Rapid insulinotropic effect of 17beta-estradiol via a plasma membrane receptor. FASEB J. 1998;12(13):1341–8.

3. Alonso-Magdalena P, Ropero AB, Carrera MP, Cederroth CR, Baquie M, Gauthier BR, et al. Pancreatic insulin content regulation by the estrogen receptor ER alpha. PLoS One. 2008;3(4):e2069.

4. Alonso-Magdalena P, Ropero AB, Garcia-Arevalo M, Soriano S, Quesada I, Muhammed SJ, et al. Antidiabetic actions of an estrogen receptor beta selective agonist. Diabetes. 2013;62(6):2015–25.

5. Tiano JP, Mauvais-Jarvis F. Importance of oestrogen receptors to preserve functional beta-cell mass in diabetes. Nat Rev Endocrinol. 2012;8(6):342–51.

6. Gannon M, Kulkarni RN, Tse HM, Mauvais-Jarvis F. Sex differences underlying pancreatic islet biology and its dysfunction. Mol Metab. 2018;15:82–91.

7. Nadal A, Alonso-Magdalena P, Soriano S, Ropero AB, Quesada I. The role of oestrogens in the adaptation of islets to insulin resistance. J Physiol. 2009;587(Pt 21):5031–7.

8. Quesada-Candela C, Tuduri E, Marroqui L, Alonso-Magdalena P, Quesada I, Nadal A. Morphological and functional adaptations of pancreatic alpha-cells during late pregnancy in the mouse. Metabolism. 2020;102:153963.

9. Wild S, Roglic G, Green A, Sicree R, King H. Global prevalence of diabetes: estimates for the year 2000 and projections for 2030. Diabetes Care. 2004;27(5):1047–53.

10. Mauvais-Jarvis F, Le May C, Tiano JP, Liu S, Kilic-Berkmen G, Kim JH. The Role of Estrogens in Pancreatic Islet Physiopathology. Adv Exp Med Biol. 2017;1043:385–99.

11. Hewitt SC, Korach KS. Estrogen Receptors: New Directions in the New Millennium. Endocr Rev. 2018;39(5):664–75.

12. Jia M, Dahlman-Wright K, Gustafsson JA. Estrogen receptor alpha and beta in health and disease. Best Pract Res Clin Endocrinol Metab. 2015;29(4):557–68.

13. Levin ER, Hammes SR. Nuclear receptors outside the nucleus: extranuclear signalling by steroid receptors. Nat Rev Mol Cell Biol. 2016;17(12):783–97.

14. Revankar CM, Cimino DF, Sklar LA, Arterburn JB, Prossnitz ER. A transmembrane intracellular estrogen receptor mediates rapid cell signaling. Science. 2005;307(5715):1625–30.

15. Thomas P, Pang Y, Filardo EJ, Dong J. Identity of an estrogen membrane receptor coupled to a G protein in human breast cancer cells. Endocrinology. 2005;146(2):624–32.

16. Wong WP, Tiano JP, Liu S, Hewitt SC, Le May C, Dalle S, et al. Extranuclear estrogen receptor-alpha stimulates NeuroD1 binding to the insulin promoter and favors insulin synthesis. Proc Natl Acad Sci U S A. 2010;107(29):13057–62.

17. Zhou Z, Ribas V, Rajbhandari P, Drew BG, Moore TM, Fluitt AH, et al. Estrogen receptor alpha protects pancreatic beta-cells from apoptosis by preserving mitochondrial function and suppressing endoplasmic reticulum stress. J Biol Chem. 2018;293(13):4735–51.

18. Le May C, Chu K, Hu M, Ortega CS, Simpson ER, Korach KS, et al. Estrogens protect pancreatic beta-cells from apoptosis and prevent insulin-deficient diabetes mellitus in mice. Proc Natl Acad Sci U S A. 2006;103(24):9232–7.

19. Soriano S, Ropero AB, Alonso-Magdalena P, Ripoll C, Quesada I, Gassner B, et al. Rapid regulation of K(ATP) channel activity by 17beta-estradiol in pancreatic beta-cells involves the estrogen receptor beta and the atrial natriuretic peptide receptor. Mol Endocrinol. 2009;23(12):1973–82.

20. Martinez-Pinna J, Marroqui L, Hmadcha A, Lopez-Beas J, Soriano S, Villar-Pazos S, et al. Oestrogen receptor beta mediates the actions of bisphenol-A on ion channel expression in mouse pancreatic beta cells. Diabetologia. 2019;62(9):1667–80.

21. Sharma G, Prossnitz ER. Mechanisms of estradiol-induced insulin secretion by the G protein-coupled estrogen receptor GPR30/GPER in pancreatic beta-cells. Endocrinology. 2011;152(8):3030–9.

22. Bian C, Bai B, Gao Q, Li S, Zhao Y. 17beta-Estradiol Regulates Glucose Metabolism and Insulin Secretion in Rat Islet beta Cells Through GPER and Akt/mTOR/GLUT2 Pathway. Front Endocrinol (Lausanne). 2019;10:531.

23. Liu S, Le May C, Wong WP, Ward RD, Clegg DJ, Marcelli M, et al. Importance of extranuclear estrogen receptor-alpha and membrane G protein-coupled estrogen receptor in pancreatic islet survival. Diabetes. 2009;58(10):2292–302.

24. Hastoy B, Godazgar M, Clark A, Nylander V, Spiliotis I, van de Bunt M, et al. Electrophysiological properties of human beta-cell lines EndoC-betaH1 and -betaH2 conform with human beta-cells. Sci Rep. 2018;8(1):16994.

25. Virgilio E, Tielens S, Bonfield G, Nian FS, Sawatani T, Vinci C, et al. Recessive TMEM167A variants cause neonatal diabetes, microcephaly, and epilepsy syndrome. J Clin Invest. 2025;135(22).

26. Costa-Junior JM, Coomans de Brachene A, Musuaya AE, Zimath PL, Martin-Vazquez E, Oliveira JG, et al. Exercise-induced meteorin-like protein protects human pancreatic beta cells from cytokine-induced apoptosis. Diabetologia. 2025;68(7):1524–9.

27. Manders EM, Stap J, Brakenhoff GJ, van Driel R, Aten JA. Dynamics of three-dimensional replication patterns during the S-phase, analysed by double labelling of DNA and confocal microscopy. J Cell Sci. 1992;103 (Pt 3):857–62.

28. Bolte S, Cordelieres FP. A guided tour into subcellular colocalization analysis in light microscopy. J Microsc. 2006;224(Pt 3):213–32.

29. Quereda C, Gomez-Vicente V, Palmero M, Martin-Nieto J. Expression of Dystroglycanopathy-Related Enzymes, POMGNT2 and POMGNT1, in the Mammalian Retina and 661W Cone-like Cell Line. Biomedicines. 2025;13(11).

30. Villar-Pazos S, Martinez-Pinna J, Castellano-Munoz M, Alonso-Magdalena P, Marroqui L, Quesada I, et al. Molecular mechanisms involved in the non-monotonic effect of bisphenol-a on ca2+ entry in mouse pancreatic beta-cells. Sci Rep. 2017;7(1):11770.

31. Rorsman P, Ashcroft FM. Pancreatic beta-Cell Electrical Activity and Insulin Secretion: Of Mice and Men. Physiol Rev. 2018;98(1):117–214.

32. Andersson LE, Valtat B, Bagge A, Sharoyko VV, Nicholls DG, Ravassard P, et al. Characterization of stimulus-secretion coupling in the human pancreatic EndoC-betaH1 beta cell line. PLoS One. 2015;10(3):e0120879.

33. Margolis KL, Bonds DE, Rodabough RJ, Tinker L, Phillips LS, Allen C, et al. Effect of oestrogen plus progestin on the incidence of diabetes in postmenopausal women: results from the Women’s Health Initiative Hormone Trial. Diabetologia. 2004;47(7):1175–87.

34. Kanaya AM, Herrington D, Vittinghoff E, Lin F, Grady D, Bittner V, et al. Glycemic effects of postmenopausal hormone therapy: the Heart and Estrogen/progestin Replacement Study. A randomized, double-blind, placebo-controlled trial. Ann Intern Med. 2003;138(1):1–9.

35. Liang C, Dobson AJ, Doust J, Ribeiro LW, Mishra GD. Menopause hormone therapy and type 2 diabetes risk and prognosis: systematic review and meta-analyses. Diabetologia. 2026.

36. Pereira RI, Casey BA, Swibas TA, Erickson CB, Wolfe P, Van Pelt RE. Timing of Estradiol Treatment After Menopause May Determine Benefit or Harm to Insulin Action. J Clin Endocrinol Metab. 2015;100(12):4456–62.

37. Finan B, Yang B, Ottaway N, Stemmer K, Muller TD, Yi CX, et al. Targeted estrogen delivery reverses the metabolic syndrome. Nat Med. 2012;18(12):1847–56.

38. Tiano JP, Tate CR, Yang BS, DiMarchi R, Mauvais-Jarvis F. Effect of targeted estrogen delivery using glucagon-like peptide-1 on insulin secretion, insulin sensitivity and glucose homeostasis. Sci Rep. 2015;5:10211.

39. Fuselier T, Mota de Sa P, Qadir MMF, Xu B, Allard C, Meyers MM, et al. Efficacy of glucagon-like peptide-1 and estrogen dual agonist in pancreatic islets protection and pre-clinical models of insulin-deficient diabetes. Cell Rep Med. 2022;3(4):100598.

40. Sanchez-Garrido MA, Serrano-Lopez V, Ruiz-Pino F, Vazquez MJ, Rodriguez-Martin A, Torres E, et al. Superior metabolic improvement of polycystic ovary syndrome traits after GLP1-based multi-agonist therapy. Nat Commun. 2024;15(1):8498.

41. Degroote L, Chrzanowski J, Lemaitre P, Wouters A, Bourgeois S, Van Mulders A, et al. GLP1-E2 therapy delays autoimmune diabetes in late-stage prediabetic NOD mice and potentiates low-dose anti-CD3 therapy for enhanced disease protection. Diabetologia. 2026;69(8):2307–22.

42. Lu W, Katzenellenbogen BS. Estrogen Receptor-beta Modulation of the ERalpha-p53 Loop Regulating Gene Expression, Proliferation, and Apoptosis in Breast Cancer. Horm Cancer. 2017;8(4):230–42.

43. Paruthiyil S, Parmar H, Kerekatte V, Cunha GR, Firestone GL, Leitman DC. Estrogen receptor beta inhibits human breast cancer cell proliferation and tumor formation by causing a G2 cell cycle arrest. Cancer Res. 2004;64(1):423–8.

44. Boronat-Belda T, Ferrero H, Al-Abdulla R, Quesada I, Gustafsson JA, Nadal A, et al. Bisphenol-A exposure during pregnancy alters pancreatic beta-cell division and mass in male mice offspring: A role for ERbeta. Food Chem Toxicol. 2020;145:111681.

